# The end of the American dream: a hard to Swallow reality - How the Barn Swallow (*Hirundo rustica*) returned from America, a complete mtDNA phylogeny

**DOI:** 10.1101/2025.06.23.661004

**Authors:** Gianluca Lombardo, Guido Roberto Gallo, Andrea De Benedictis, Marta Cavallini, Giulio Formenti, Roberto Ambrosini, Luca Gianfranceschi, Giorgio Binelli

## Abstract

The barn swallow (*Hirundo rustica, H.r.*) is one of the most iconic and fascinating migratory birds since ancient times, their philopatry and pair bonding have been the symbol of travellers and lovers throughout history. The barn swallow subspecies complex has been the main focus of many latest papers, focusing mainly on the Eurasian and Levant subspecies (*H.r. rustica* and *H.r. transitiva*). The remaining subspecies have either not been studied or few complete mitogenome sequences have been obtained. We present a comprehensive phylogeographic analysis of 580 complete mitochondrial genomes of barn swallows representing all recognized subspecies. We identified 553 unique haplotypes, 166 of which are novel, with high haplotype diversity and notable nucleotide diversity among subspecies. Phylogenetic analysis by maximum parsimony supports a single maternal ancestor and reveals five main haplogroups, confirming and refining prior classifications. A novel, distinct haplogroup (E) was identified for *H.r. tytleri*, previously classified as a sub-branch of *H.r. erythrogaster*. The inclusion of 66 hybrid individuals confirmed extensive maternal introgression and revealed multiple cases of mitogenome-subspecies mismatches, notably in *savignii*, *transitiva*, and *tytleri* populations. We further integrated 13 mitogenomes from 12 other *Hirundo* species, three of which are newly sequenced, to contextualize *H. rustica* evolution within the genus. Our findings support a southern African origin with subsequent diversification and a northward expansion. This study significantly enhances resolution of barn swallow mitogenomic diversity, revises phylogenetic relationships, identifies new haplogroups, hints at inter-subspecies gene flow and refines coalescent-based divergence times, offering key insights into the evolutionary history and hybridisation patterns of this iconic migratory species.

## Introduction

The iconic barn swallow (*Hirundo rustica*) stands out of the flocks as one of the most extensively distributed avian species in the world (Turner and Rose 2010). This phenomenon is mostly attributed to its social behaviour regarding nesting, switching from natural nesting sites like caves and cliffs, to human-made structures (Zink et al. 2006). The species is commensal with humans and its history is intertwined with our culture as well, we can in fact see swallows appear in art, poetry, legends and myths of different cultures (Green 2019). The *H. rustica* complex comprises at least five species: *Hirundo rustica rustica* (*H.r. rustica*) breeding in Eurasia and North Africa; *H.r. savignii* a non-migratory resident of Egypt; *H.r. gutturalis* breeding in Japan, Korea, China, Mongolia and Eastern Russia; *H.r. tytleri* breeding in Russia on the border with Mongolia and China and *H.r. erythrogaster* breeding from Alaska to Mexico. Another population has been historically defined in Israel as the resident *H.r. transitiva* subspecies. Taxonomy has been based traditionally on both morphology, having different coloured plumage, breastbands, size and tail lengths and its behaviour and given its non-migratory status. However, recent studies have shown that the genetic difference in mitochondrial DNAs (mtDNA) within *transitiva* and between *rustica* are of the same magnitude, indicating they are the same subspecies (Dor et al. 2012; Lombardo et al. 2022), at least from the mitochondrial point of view.

Moreover, hybridisation zones have been described and involve all subspecies (from here on just indicated by subspecies name) except for the American *erythrogaster* who are isolated in the Americas. In the Eastern Mediterranean region, the migratory routes of *rustica* overlap with those of *savignii* and *transitiva*, leading to genetic admixture such as *savignii* mitochondrial DNA found in phenotypical *transitiva* samples and *rustica* mitogenomes found in *savignii* samples and *transitiva* (Dor et al. 2010; Dor et al. 2012; Carter et al. 2020; Lombardo et al. 2022). On the other hand, in the East migratory divides exist between: *rustica*-*tytleri* in the Siberian Oblast of Irkutsk; *rustica*-*gutturalis* in the Qinghai and Gansu regions of China and *tytleri-gutturalis* in NE-Mongolia and Buryatia-Zabaykalsk in Russia (Scordato et al. 2017; Liu et al. 2020; Lombardo et al. 2022).

Advances in genomic technologies — particularly whole genome sequencing (WGS), shotgun sequencing, and genotyping-by-sequencing — have dramatically increased the amount and the availability of never-before-seen genetic data. However, most mitochondrial reads from WGS datasets remain underutilized, while harbouring great potential for phylogenetic analyses, especially in unmapped mitochondrial reads. To date, genomic studies of barn swallows have primarily focused on nuclear genome structure, including the first species pangenome (Formenti et al. 2019; Secomandi et al. 2023), and on population history and admixture (Smith et al. 2018). These efforts enabled a deeper exploration of the conservation and evolutionary dynamics within the barn swallow genome — especially for *rustica*, *savignii* and *erythrogaster* — leading to the identification of a comprehensive set of genetic markers and candidate genomic regions under selection.

Both mitochondrial and nuclear data support the species’ monophyly and identify two major phylogenetic clusters, these being 1. *rustica* (and *transitiva*) with *savignii* and 2. *gutturalis* with *erythrogaster* and *tytleri* (Sheldon et al. 2005; Zink et al. 2006; Dor et al. 2010; Dor et al. 2012; Malaitad et al. 2016; Smith et al. 2018; Lombardo et al. 2022). The origin of an ancestral barn swallow was dated to ∼277 thousand years ago (kya). Despite ongoing sequencing efforts, data on *gutturalis* and *tytleri* subspecies remain minimal, particularly regarding complete mitochondrial genomes, resulting in imprecise divergence times estimates. For example, *tytleri*’s expansion into the Baikal region from North America has been dated between ∼20 kya (Lombardo et al. 2022) and ∼27 kya (Zink et al. 2006), based on mitogenome coding regions and the complete mitochondrial *ND2* gene, respectively.

In this study, we leverage publicly available whole genome sequences to reconstruct complete mitogenomes from under-sampled barn swallow subspecies and refine coalescent-based divergence time estimates to improve phylogenetic resolution.

## Results

### Subspecies Genetic Diversity in Barn Swallows

In silico extraction of mitochondrial reads was employed to retrieve 168 novel *Hirundo rustica* mitogenomes belonging to all postulated subspecies spanning the entire breeding range. Sequences taken from GenBank were all from whole genome studies and were therefore not enriched for mitochondrial reads. On average initial raw fastq files were ∼8000 reads, after the removal of duplicates and adapters, and mapping of reads we measured an average of 15% mitochondrial reads still present. These were enough to cover, on average, 93% of the entire mtDNA sequence with a mean coverage of 30X. GC count was 44.9%. Coding regions (nps: 1-14860; 16068-16741; 18075-18143) of new barn swallow samples revealed 166 new haplotypes (Hd = 0.99). When considering the complete dataset, a total of 553 unique haplotypes were found (Haplotype diversity, Hd = 0.95) with 1956 polymorphic sites in the coding region and, 2039 different mutations of which, 954 were private mutations. Average coding region (15,601 bp) nucleotide diversity (π) for all *Hirundo rustica* individuals is 0.543% (±0.023%), the largest variability was within the *erythrogaster* subspecies, π = 0.195% (±0.024%) and lowest in *savignii,* π = 0.076% (±0.008%) (Table 1).

Control regions were then analysed and the mutational patterns in Table 2 were found for all main haplogroups. These were added into previously published the mtPhyl script (Lombardo et al. 2022) to obtain a more detailed picture of sub-haplogroup specificity. Control region pairwise identity was 86% (1149 bp out of 1271 bp) identical and therefore mutations usually mapped twice, once in the CR1 and once in the repeated portion of the CR2. In addition, the number of repeats in the CR2 were uncountable due to the limitations of NGS and therefore were aligned to the reference sequence and were not used for phylogeny.

### Hirundo rustica tytleri, a new haplogroup in the barn swallow phylogeny

Using Bayesian tip dating molecular techniques, we obtained coalescence times for main divergence events for all five main haplogroups. Haplogroup A (*N* = 389) expanded in Eurasia around 64 ±0.7 kya (Table S1). The four main haplogroups of A1a1-A1a4 (Figure S2) all diverged at the beginning of the LGM 23-33 ± 0.1 kya (Figure 1). Haplogroup B (*N* = 12) dated 24 ± 0.3 kya is subdivided into three main sub-haplogroups, B1 in Israel (already previously described (Lombardo et al. 2022)) and B2 and B3 in Egypt (Figure S3). These diverged respectively 11, 19 and 3.6 (±0.1) kya. Haplogroup C originated 40 ± 0.3 kya and split into two main haplogroups (Figure S4), C1 and C2 both encompassing samples from Russia, China, Mongolia and Japan indistinctively. Haplogroup C1, dates back to 32 ± 0.2 kya and further differentiates into three sub-haplogroups, C1a-c, once again without geo-specificity (20-30 ± 0.2 kya). Haplogroup C2 originated 35 ± 0.4 kya and splits into five sub-haplogroups, C2a-C2e, C2b and C2e are Mongolia (Ugii Lake area) and China-specific (E-SE China) respectively (13-27 ± ∼0.2 kya). Other interesting geo-specific sub-haplogroups are found closer to the leaves of the phylogenetic tree, these are: C1c1a, Northern/ Eastern China and C2d1a1 circumscribing the Baikal Lake (Southern Siberia) and dated 17 ± 0.1 kya. Haplogroup D (*N* = 24), now dated to 66 ± 0.7 kya (named DE in Table S1) and is now characterised by three main clades (Figure S5), D1, D2 and E, more on the latter haplogroup later. Both main haplogroups are also older than previously thought, D1, 35 ± 0.7 kya and D2, 30 ± 0.2 kya. Main sub-haplogroups are now: D1a (USA and Canada), D2a (Nebraska and Colorado) and D2b (Nebraska) (20-24 ± 0.2 kya). This new haplogroup E diverged from the American subspecies around 58 ± 0.6 kya and split into two main sub-haplogroups (Figure S6), E1 (33 ± 0.2 kya) and E2 (37 ± 0.5 kya) both containing Mongolian and Russian birds. Notably a Chinese bird registered as *rustica-gutturalis* hybrid present in sub-haplogroup E1a1b and an American one in E2a.

**Figure 1:**
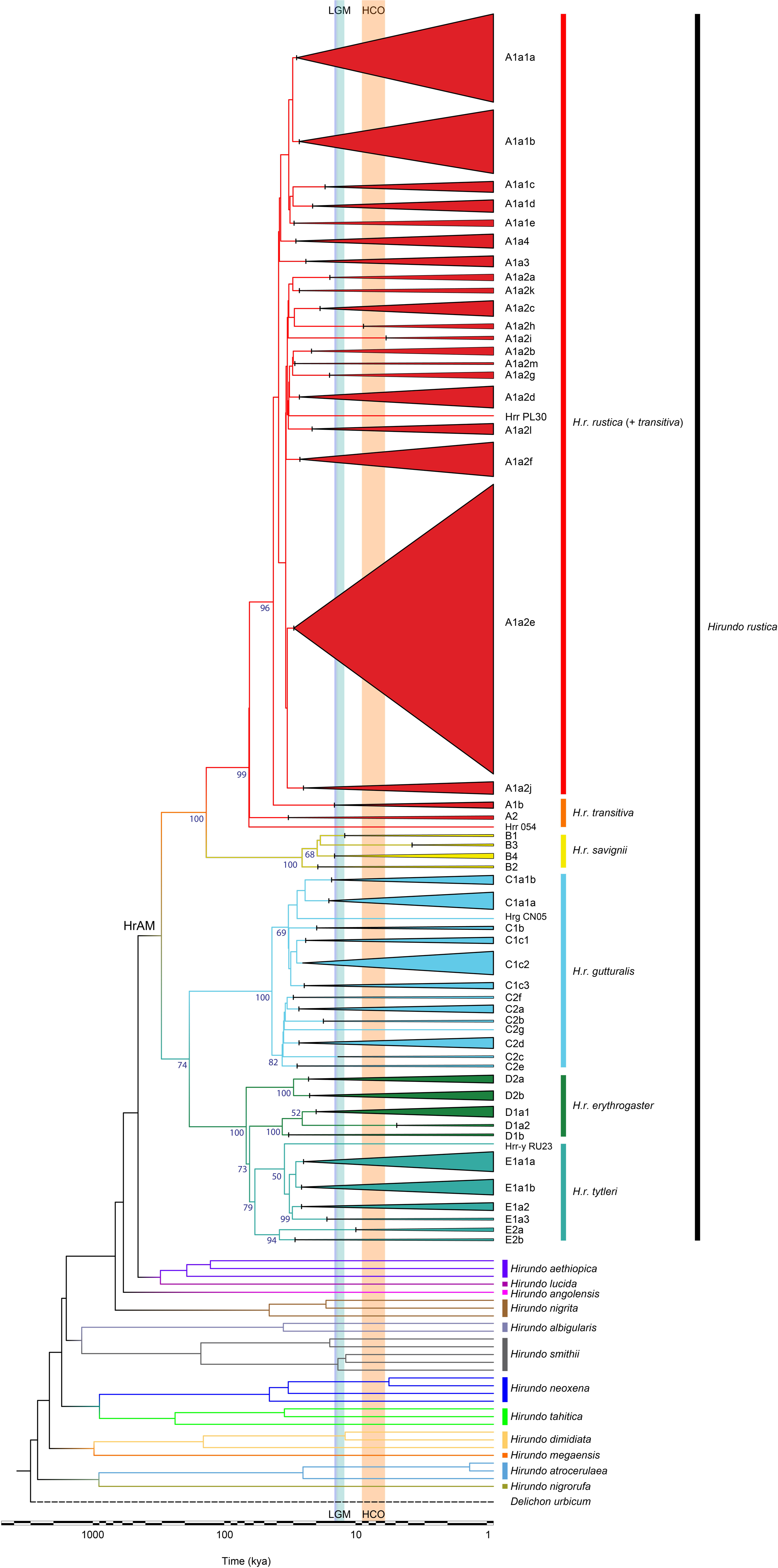
Schematic MP phylogeny of Hirundo rustica mitogenomes and closest species. This tree was built using the entire mitogenome coding-region (15,601 bps; nps 1-14,859, nps 16,068-16740, nps 18,075-18,143) of 580 barn swallows and 30 individuals of the closest species (all Hirundo). The tree was rooted using Delichon urbicum (NC_050298). Major sub-haplogroups of H. rustica are represented as triangles which are proportional to the number of mtDNAs present. HrAM refers to the Hirundo rustica Ancestral Mitogenome. Colours were assigned based on subspecies for the barn swal­ low while the other colours indicate the other species. Bootstrap values (1000 iterations) are shown for major haplogroups of H. rustica. The timeline (log1O) at the bottom refers to the Bayesian coalescence times of Supplementary Table S1.

### The complete Phylogeographical history of Swallows

We calculated the ancestral barn swallow mitogenome progenitor to have lived 302 ± 2 kya (Figure 2). From this ancestral population two main clades diverged. The oldest main calde is that leading to the eastern and American populations (CDE), occurring 167 ± 3 kya. Then the ancestor leading to Eurasia and Egypt (AB) diverged 128 ± 2 kya. Node AB further splits off into haplogroup A (comprising both *rustica* and *transitiva*) 64 ± 0.7 kya and haplogroup B, 24 ± 0.3 kya, both dates are slightly older than previously calculated due to the generally older split calculated for HrAM.

**Figure 2.**
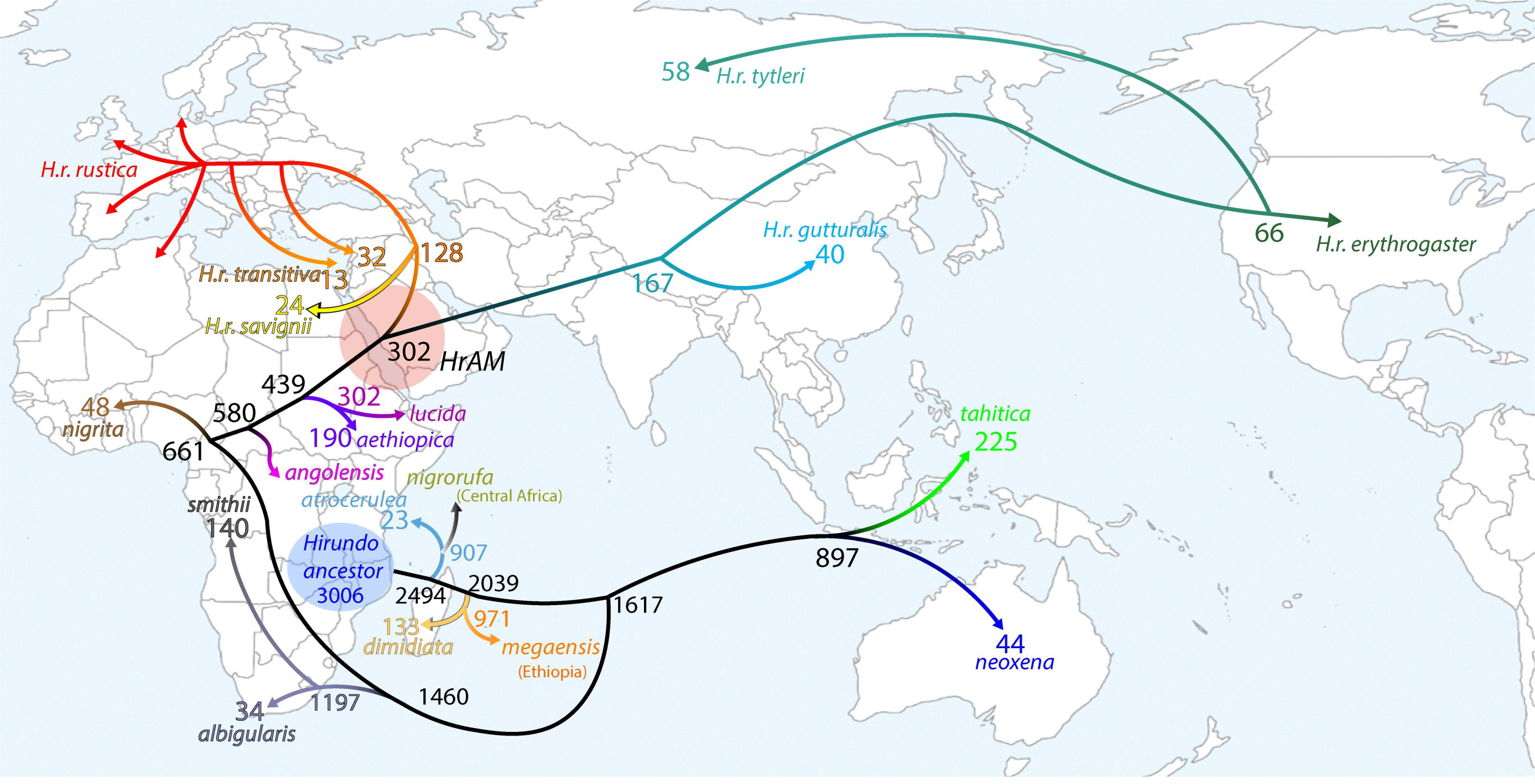
A model for the geographical and temporal spread of swallows. Map showing time divergence and hypothetical splits and diffusion routes of Hirundo species and barn swallow haplogroups. The dashed grey circle indicates the possible homeland of the Hirundo rustica mitochondrial Eve (HrAM), while the other dashed circles indicate zones where two subspecies are currently found (RT, rustica-tytleri; TG, tytleri-gutturalis and RG, rustica-gutturalis), indicating recent admixture between subspecies. Note that haplogroup A arrows are not related to any specific haplogroup but are just there to show the vast expansion of the subspecies.

We obtained 13 additional mitogenome coding region belonging to twelve different *Hirundo* species. Their phylogeny roughly follows a South-to-North pattern. Three of these have never been published before (*H. nigrorufa*, *H. megaensis and H. lucid*a). We observed *H. aethiopica* and *H. lucida* are sister groups still present in the putative origin area of HrAM and are in fact at the base of the *Hirundo rustica* monophyletic clade which separated 302 ± 8 kya. Modern *H. aethiopica* haplogroups are dated to 190 ± 6 kya. *H. angolensis* is at the base of the aforementioned groups unlike previously seen (Sheldon et al. 2005; Dor et al. 2010) and split around 580 ± 19 kya, followed by *H. nigrita* which split earlier (661 ± 20 kya) and expanded West with current mitogenomes having a 48 ± 2 kya common origin. *H. albigularis* and *H. smithii* form a monophyletic sister clade which split from the main phylogeny in southern Africa around 1462 ± 60 kya, the two species then diversified from one another around 1197 ± 31 kya with *albigularis* remaining in southern Africa (34 ± 1 kya) while *H. smithii* spread throughout the continent in two major haplogroups (SMI-A, 14 ± 0.2 kya and SMI-B, 13 ± 0.1 kya). We then see the major split which occurred 1617 ± 53 kya between African swallows and the Pacific clade which comprises *H. tahitica* and *H. neoxena*. These diverged from each other 897 ± 55 kya and reached the Australasian and Oriental areas with current ancestries originating respectively 225 ± 16 kya and 44 ± 0.8 kya. The second-to-last basal phylogenetic split occurred in South-East Africa 2039 ± 70 kya, and gave rise to the monophyletic sister clade comprising *H. dimidiata* and *H. megaensis*, which had been previously described as the “Pearl-breasted clade”. These two diverged 971 ± 54 kya with the latter reaching the Ethiopian state of Oromia, *H. dimidiata* instead remained in southern Africa and its origins can be traced back to 133 ± 8 kya. Finally, the most basal clade split from the hypothetical *Hirundo* ancestor 2494 ± 65 kya and encompasses *H. nigrorufa* and *H. atrocerulea* also known as the blue clade. This clade originated 907 ± 45 kya with both species remaining in Central Africa. The *H. atrocerulea* ancestor can be traced back recently to 23 ± 0.5 kya (based on 3 modern mitogenomes).

## Discussion

### Swallow Fingerprinting and species distribution

The greatest hurdle in mapping the complete mitochondrial DNA in swallows is the duplicated control region, also found in many avian species (Urantówka et al. 2020) and especially in passerines (Mackiewicz et al. 2019). In our case the duplicated control region reads are indistinguishable from one another when not considering the 80 base pairs which are different in the CR2 and the final 210 bps which are a long tandem repeat in CR2. This occurred for every individual and therefore were included just for phylogeny and not for dating branches as double the mutations in the CR regions would increment divergence times erroneously. Moreover, we uncovered a pattern in the CR mutations that are subspecies specific and could be used to develop a cheap molecular method for subspecies determination that does not rely on phenotype or geography, useful for hybrid zones. GenBank and ENA were also scoured for *H. rustica* mtDNA fragments which can be used to produce a barn swallow SNP dataset to fingerprint samples at the subspecies level, or even better, at the sub-haplogroup level. These samples were from Project PRJNA323498, mainly 100bp single-end reads previously digested with Msel and EcoRI which maps on the mtDNA of most samples at rRNA 12s (829-907), *ND2* (4839-4923; 4924-4990), *COI* (5866-5931; 6215-6300), *COII* (7155-7302; 7310-7394), *ATP6* (8394-8459), *ND4* (11166-11251; 11259-11342), *cytb* (14251-14335), CR1 (15140-15224; 15232-15315) and CR2 (17045-17129; 17137-17220). This made it possible to determine 89.3% of the samples, or 1280 individuals at least to the subspecies level (Figure 3). This led to the discovery of two additional sub-haplogroup of *H. savignii* named B4 and B5 where a rather large number (N=27) of mitogenomes cluster in. This indicates the unsampled potential in genetic variability in the subspecies breeding range. All other samples cluster into previously described major sub-haplogroups. Note that all these partial sequences were only used for internal classification and not for any phylogenetic or temporal analysis. Noteworthy individuals were two *erythrogaster* samples with *tytleri* mitogenomes present in Colorado, supporting our title’s claim, six *gutturalis* samples presenting either *rustica* or *tytleri* mitogenomes, six rustica samples presenting either *gutturalis* or *tytleri* mitogenomes and six tytleri with gutturalis mitogenomes all sampled in the hybrid zones. Moreover, two *savignii* samples with *rustica* mitogenomes and four *transitiva* with *savignii* mitogenomes confirming the region as another hybrid zone. These samples were useful to uncover the maternal lineage of hybrid individuals in particular. In fact, 486 individuals were hybrids and 463 (95%) could be identified at the subspecies level. Of the 683 single or multi gene accessions, 671 (98%) could be identified to the subspecies level indicating that the resolution of our phylogeny is well structured enough to distinguish between subspecies using few SNPS.

**Figure 3.**
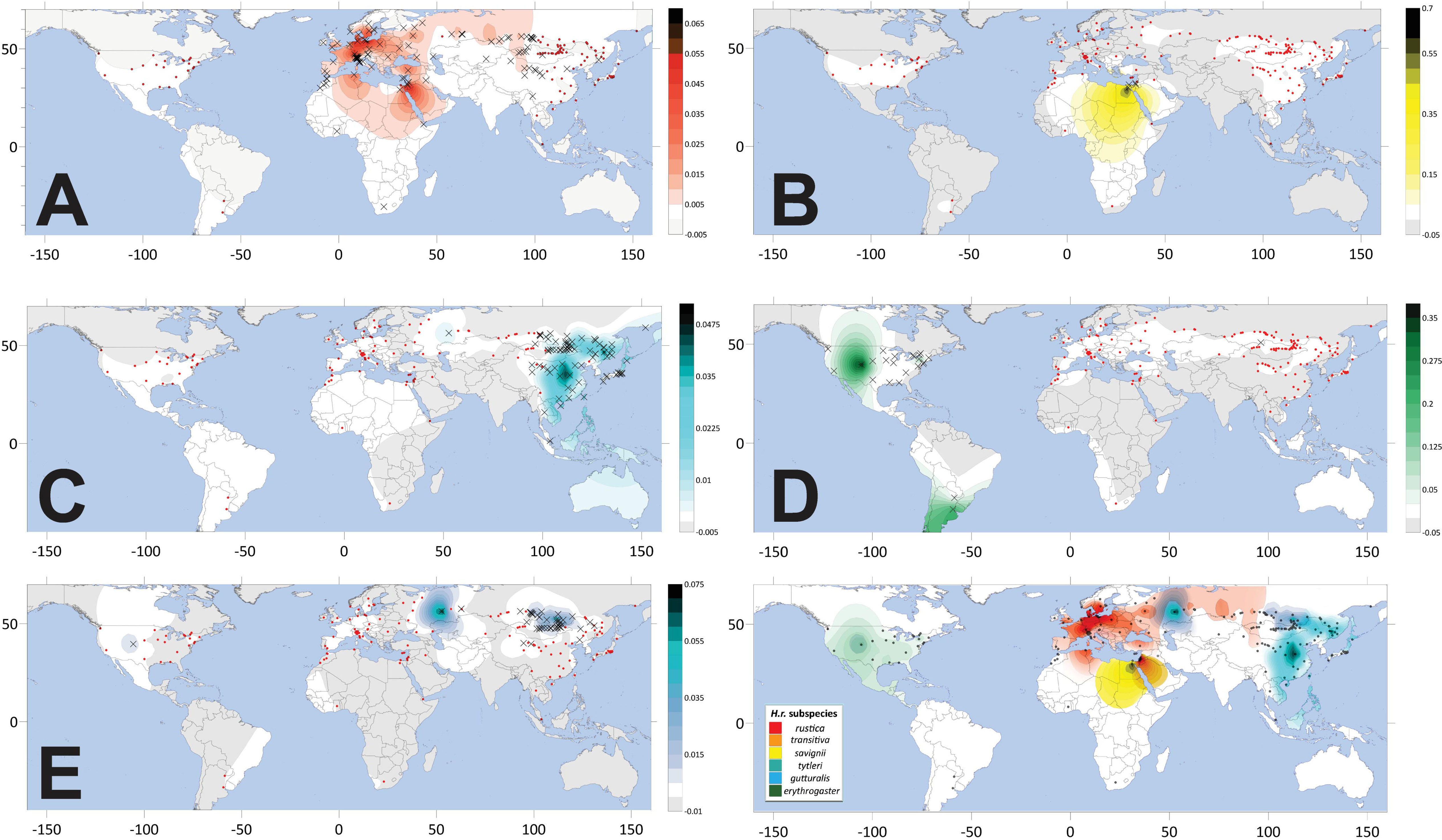
Haplogroup frequency distribution maps of main barn swallow haplogroups (A-E). Dots indicate the geographical locations of all sampled individuals; Crosses indicate individuals of subspecies in question. Colour scale indicates frequency of haplogroup in the given map. The final map is a composite of all subspecies distribution breeding ranges of all sampled individuals and database individuals.

### Hirundo rustica, a richer phylogeny

Phylogenetic analysis of the 580 complete mitogenomes coding region (15,601 bp) revealed the species is monophyletic deriving from a single ancestral female (HrAM). The addition of the new samples (Figure 4) maintain the same grand phylogenetic structure as previous findings (Zink et al. 2006; Dor et al. 2010; Lombardo et al. 2022), showing two major clades splitting from the origin encompassing haplogroups A and B in one and C and D in the other. A major overhaul in haplotype classification can be seen for haplogroup C (*N* = 72), for simplicity, the previously published four samples have been re-classified according to the new nomenclature here described maintaining however the two main sub-haplogroups structure. One new feature of crucial importance in the phylogenetic tree which appears in this study is the new “main” haplogroup E, in fact, most *H.r. tytleri* were classified as a sub-haplogroup of the American *H.r. erythrogaster* (haplogroup D) given the single SNP that defines the branch. That said, the clade was distinct enough from the others, with π = 0.285% (±0.015%), so has been named haplogroup E for simplicity’s sake (and not D3 as previously theorised from cytb sequences, Lombardo et al., 2022).

**Figure 4.**
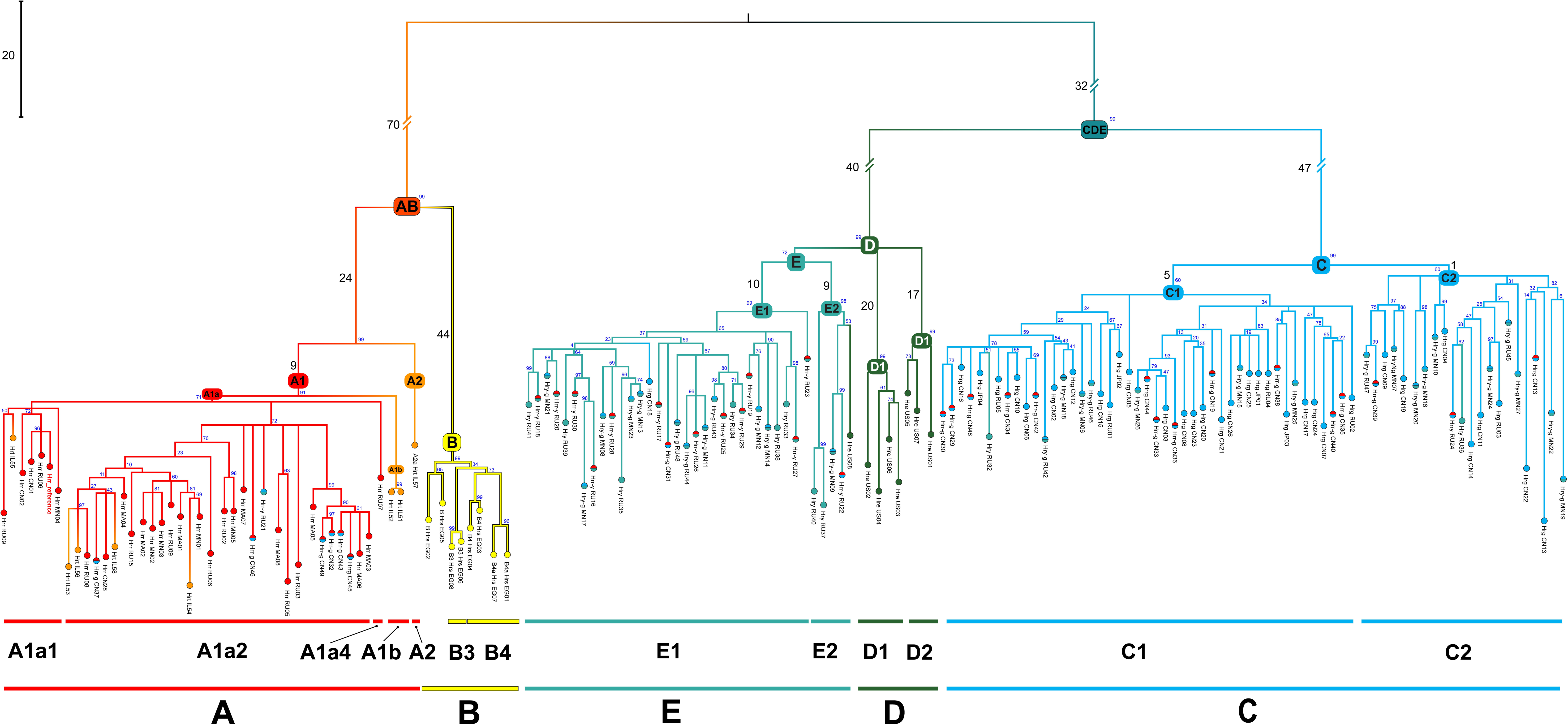
Maximum Parsimony tree of Hirundo rustica mitogenomes. This tree encompasses only new complete samples added to the study for clarity. Colours indicate main subspecies. Numbers on main branches indicate number of nucleotide substitutions while numbers at main nodes indicate bootstrap values.

### An insight into transitiva

When analysing swallow samples from the Levant region we found Haplogroup B. Having been previously defined by only four entire mitogenomes from Israel, it could not be said with 100% certainty, at least from the complete mtDNA point of view, that Egyptian *savignii* harboured the same DNA as the four *transitiva* (based on morphology and behaviour) already present in this haplogroup, this was in fact only postulated using *cytb* sequences (Dor et al. 2010; Lombardo et al. 2022). The addition of eight complete *savignii* sequences (sampled in Egypt with correct phenotypic traits) (Smith et al. 2018) confirms the four noted Haplogroup B individuals from Israel are therefore present in the previously postulated hybrid zone in the Sinai Peninsula, a genetic and geographic crossroad between the three subspecies in the area. All *transitiva* individuals added in this study are all found within haplogroup A as previously seen. The behaviour and phenotype of *H.r. transitiva* being non-migratory with a rufous breast with reddish underparts respectively are more similar to *savignii* than *rustica*, this indicates an intermediate phenotype between *rustica* and *savignii*. Moreover, a curious case can be made for haplotypes A1b and A2 (Figure 5). These are mainly *transitiva* individuals with the exception of a Polish and Algerian *rustica* (A1b) and a Spanish and Italian *rustica* (A2). Both these haplogroups have a rather noteworthy number of defining private mutations (11 and 17 respectively) which indicate an older ancestral origin of these haplogroups with respect to other rustica haplogroups, this can be a hint towards a subspeciation event (or two) (nucleotide diversity values, π ≈ 0.2-0.3%).

**Figure 5.**
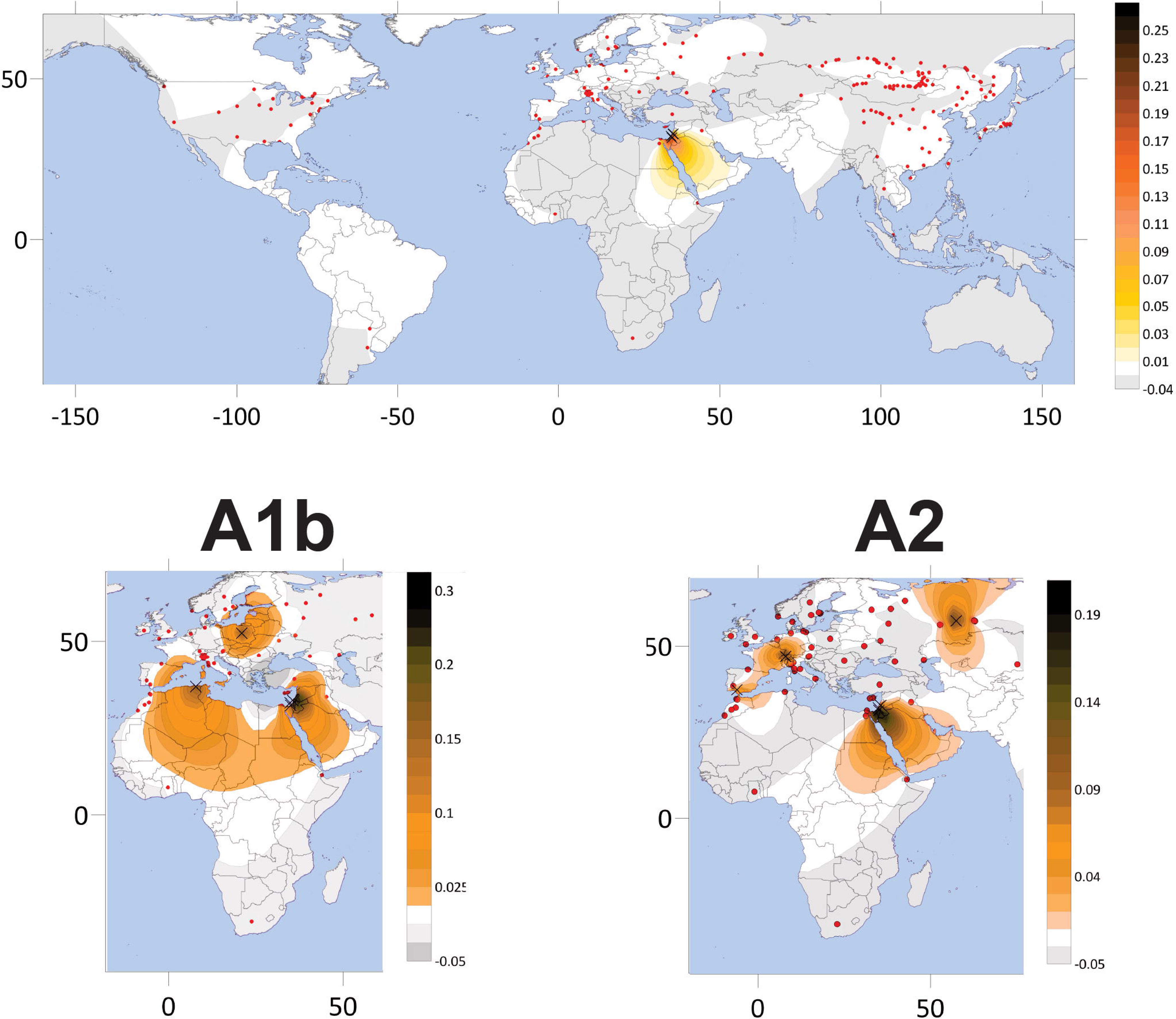
Haplogroup frequency distribution maps of H.r. transitiva individuals. Dots indicate the geographical locations of all sampled individuals; Crosses indicate individuals of subspecies in question. Colour scale indicates frequency of haplogroup in the given map.

Our obtained nucleotide diversity values are in the range of those seen for other species of birds and their subspecies (Questiau et al. 1998; White et al. 2013; Lombardo et al. 2022), with A1b being on the lower end at π = 0.175% and on the other hand A2 could be more in line with a subspecies π = 0.294%. Furthermore, five samples from Israel identified via phenotypical characteristics (Dor et al. 2012, Smith et al. 2018) and *cytb* and CR fragments cluster within haplogroups A1b and A2 lending further credence to this hypothesis. What’s more, A2 diverged pre-LGM and is more likely the ancestral *transitiva* mitogenome. This lineage either stayed in the Levant region or migrated south during the LGM and returned post Younger Dryas until present day as seen in other migratory birds (Zink & Gardner 2017; Carrera et al., 2022; Kimmit et al., 2023; Ferrer Obiol et al., 2025). This pattern can be seen in the population structure analysis of nuclear genomes where *transitiva* individuals have three patterns corresponding to the two hybrids and a “pure” *transitiva* lineage (Gallo et al. in preparation). Geneflow still occurs between migrating *rustica* and *transitiva* individuals and therefore either have not had enough isolation to properly diversify, or as can be seen by the four haplogroup B *transitiva* individuals, haplogroup A replaced the previously present haplogroup B (still present in Egypt).

### The end of the American dream

The trans-Beringian colonization direction of *H.r. tytleri* goes against the current of many described Nearctic-palearctic taxa (Kurt & Cook, 2004) and has been postulated to have occurred with the arrival of humans and their artifacts in the area (Smirenskiy & Mishchenko, 1981). Given our calculations this happened in two main dispersion events, those of haplogroup E1 and E2. These diverge from a *Hirundo rustica erythrogaster* ancestor which back migrated around 33-37 kya, in line with what previously described in another study based on few ND2 genes (Zink et al., 2006). Haplogroups E1 and E2 share a single SNP in the coding region in common at the E root, calculated to have diverged around 58 kya and have greatly diverged since without geo-specificity as both are found throughout the distribution range. Interestingly we have an American (*H.r. erythrogaster*) individual that carries a *tytleri* mitogenome, this is a further proof of the American exit as this mitogenome remained in the American continent and gained mutations alongside its sister clade E2a1. Given the considerable number of ND2 and COI mutations on the DE node (7 and 5 respectively), even partial sequences present in the database could be clearly identified (Not shown in this work) to the sub-haplogroup level (albeit could not be used for time estimations)

### Hirundo rustica hybrids

Unlike the migratory divide between *rustica-gutturalis* in Western China, and that of *tytleri-gutturalis* in Northern Mongolia and the border between Russia and China, the hybrid samples from the Irkutsk Oblast region of Russia (*rustica-tytleri* hybrids) show a significant skew from the expected haplogroup distribution (Figure 6). These have a majority of *tytleri* mitogenomes indicating that maternal lineages in the hybrid regions are mostly *tytleri*. This would seem to suggest a sign of sexual selection preference of female *tytleri* for male *rustica* individuals as also seen by the hybrid phenotypes which are intermediate, at the contact zones which goes against what has been previously studied using genomic data where patterns of sexual selection as reproductive barriers were uncovered (Vortman et al., 2011; Schield et al., 2024), at least, just for *rustica-tytleri* and only from the mitogenome point of view.

### Before the Barn: Uncovering the Swallow’s Past

We obtained new and updated ages for all nodes in the phylogeny using Bayesian age estimates (Table S1). With the expansion of the previous major haplogroups and addition of the new Haplogroup E we have found older nodes present in the East thus, pre-dating the ancestral barn swallow mitogenome progenitor (HrAM) to 302 ± 8 thousands of years ago, close to the higher SD of the previous study (Lombardo et al. 2022). This event could have taken place in North-eastern Africa (Sudan/Ethiopia/Somalia) as the closest species (*H. ethiopica* and *H. lucida*) are still present there in current day (Figure 2) (BirdLife International et al. 2016A; BirdLife International et al. 2017). Moreover, given the extensive phylogeny in this study, we traceback the origin of all *Hirundo* species to a common mitochondrial ancestor 2.4 Mya in what was probably Southern Africa, given that the most basal species are still present there. From here the phylogeny follows a South-to-North patten of colonisation with most recent species being more north than species which split before, arriving at worldwide expansion of *H. rustica*. It is to note that the Pacific clade branched off 1.6 Mya (in line with what can be seen from nuclear data, Broyles et al. 2023) and reached southeast Asia and Australia vastly before *H. rustica* expanded. This most likely occurred via island hopping (Broyles et al. 2023; Estandía & Recalde, 2024). The three new complete mitogenomes, *H. nigrorufa*, *H. megaensis* and *H. lucida* fall into previously described clades with other species as in previous *ND2* and *cytb* studies. One major change to the phylogeny is the placement of *H. albigularis* which was previously basal to the pacific clade and the *Hirundo* clade (Dor et al. 2010), given incomplete data, and is now a sister species of *H. smithii.* The latter is another interesting species as two main subspecies have been described, *H.s. smithii* and *H.s. filifera*.

One present throughout Africa and the other present in Asia (BirdLife International et al. 2016B). We see two major haplogroups present which separated from each other around 140 kya, dates comparable to the subspeciation of *H. rustica*. These are indicative of the presence of ancestral African mitochondrial DNAs belonging to the Asian subspecies.

## Conclusions

This study presents the most extensive mitogenomic analysis of the barn swallow (*Hirundo rustica*) to date, encompassing 580 high-quality mitochondrial genomes across all six recognized subspecies. Our results reveal 553 unique haplotypes, underscoring the substantial intraspecific diversity of this species, and demonstrate clear phylogeographic structure through five well-supported haplogroups. The discovery of haplogroup E, largely restricted to *H.r. tytleri*, adds complexity to the mitochondrial landscape of *H. rustica*, highlighting previously underappreciated evolutionary lineages. Furthermore, we argue that *H.r. transitiva*, though previously discounted as a subspecies, could actually be considered one given two geo-specific sub-haplogroups of A. Time-calibrated phylogenies place the most recent common ancestor of *H. rustica* mitogenomes at ∼302 kya, with divergence among major haplogroups between 128–47 kya, likely reflecting responses to Late Pleistocene climatic oscillations. Patterns of haplotype diversity and demographic reconstructions support scenarios of historical expansions and secondary contacts, particularly in the Palearctic region. Signals of mitochondrial introgression in hybrid zones involving mainly, *H.r. transitiva* in the Sinai Peninsula, and *H.r. tytleri* in northern Mongolia align with previous nuclear genome evidence, suggesting recurrent gene flow and incomplete lineage sorting. In contrast, subspecies such as *H.r. erythrogaster* maintain high diversity and distinct mitochondrial signatures, indicative of long-term demographic stability. The phylogeny of the genus *Hirundo*, constructed from mitogenomes of 13 species, supports the postulated northeastern African origin of *H. rustica* and reinforces its placement within a clade of long-distance Afro-Palearctic migrants. Additionally, analysis of the mitochondrial control region revealed lineage-specific structural variants, offering a powerful tool for high-resolution, low-cost maternal lineage identification. These findings not only resolve key aspects of the evolutionary history of *H. rustica* but also lay a foundation for future research. Integrative studies combining mitogenomic and nuclear data will be essential to disentangle historical demography from selective and stochastic processes shaping current diversity. Expanded sampling from undersampled regions, especially in Central Asia, Northern Africa and North America will further refine biogeographic and demographic models. Moreover, the control region markers identified here hold promise for conservation monitoring and for linking mitolineages to phenotypic, behavioral, or ecological variation under natural and anthropogenic pressures.

Overall, this work contributes a robust mitogenomic framework for a model migratory passerine and offers new insights into the evolutionary dynamics of widespread, phenotypically diverse species.

## Materials and Methods

### Samples Analysed

We obtained a total of 168 barn swallow mitogenomes by extracting mitochondrial reads from genomic Bioprojects (PRJNA304409, PRJNA323498, PRJNA545868, PRJNA909772, PRJNA1117501) (Smith et al. 2018; Oliveros et al. 2019; Feng et al. 2020; Secomandi et al. 2023; Schield et al. 2024). These include individuals from six subspecies: 30 *H.r. rustica* from Europe, Asia and North Africa; 8 *H.r. savignii* from Egypt; 8 *H.r. transitiva* from Israel; 35 *H.r. gutturalis* from Russia, China and Japan; 10 *H.r. erythrogaster* from the U.S.A. and 11 *H.r. tytleri* from Russia. Moreover, mitogenomes from three hybrid populations were obtained: 21 *rustica-gutturalis* hybrids from China; 16 *rustica-tytleri* hybrids from Russia and 29 t*ytleri-gutturalis* hybrids from Mongolia and Russia (Table S3). In addition to barn swallow subspecies, we extracted and obtained the mitogenomes of 12 other *Hirundo* species (*H. atrocaerulea, H. nigrorufa, H. megaensis, H. dimidiata, H. tahitica, H. neoxena, H. albigularis, H. smithii, H. nigrita, H. angolensis, H. lucida, H. aethiopica*). Remaining sequences were downloaded from Genbank, these include 411 samples from Bioproject (PRJEB51610) (Lombardo et al. 2022) being: 342 *H.r. rustica*, 50 *H.r. transitiva* (of which 4 harbouring *H.r. savignii* mitogenome), 5 H.r. gutturalis (1 harbouring *H.r. rustica* mitogenome) and 15 *H.r. erythrogaster*. Other deposited complete mitogenomes were found to be erroneous in a previous work (Lombardo et al. 2022) and were not included, aside from KX398931 (*H.r. erythrogaster*, Keepers et al. 2016, direct submission) whose small NUMT was considered as a missing segment. Furthermore, for fingerprinting analysis, we downloaded 1440 fastq files belonging to Bioproject PRJNA323498 (Smith et al. 2018) and 1509 accessions of barn swallow genes belonging to 683 different individuals. These were then aligned to the *H.r. rustica* reference sequence (MZ905359) (Lombardo 2021) using the fast Geneious mapper. Variant calling was performed using the Geneious algorithm and samples were classified by hand by comparing individuals to the complete mtDNA alignment.

### Mitochondrial read extraction and mapping

Mitogenome reads were extracted from genomic reads using an in-house pipeline. Data was downloaded from the sequence read archive (SRA) by filtering for the term *Hirundo* as paired end fastq files, NGS long reads or complete genomic scaffolds. The raw single file samples were firstly loaded into Geneious Prime v.2024.0.5 (https://www.geneious.com, Dotmatics, Boston MA) and filtered by removing duplicate sequences using Dedupe from BBTools, then aligned and mapped to the barn swallow reference sequence (MZ905359)(Lombardo 2021) using the Geneious mapper. On the other hand, samples with paired end sequences were run through an in-house pipeline for swallow analysis. Adapters and low-quality ends with phred score below 30 (99.9% likely) were removed with TrimGalore v.0.6.10 (Krueger et al. 2023), proper adapter flags were set for each dataset. Trimming efficiency was then checked using FastQC v.12.0 (Andrews 2010) to determine quality of remaining reads. Files were then aligned and mapped to the barn swallow reference sequence (MZ905359) using BWA-MEM v. 0.7.17 (Li 2013). Resulting BAM files were uploaded to Geneious Prime for manual curation. Variant calling was performed using the Geneious algorithm when at least 70% of reads shared a mutation, otherwise they were considered heteroplasmies. An exception was made for node defining mutations when they were expected. Consensus for each file was then output as a *fasta* file and aligned with Genbank entries using MEGAX v.11.0.11 (Kumar et al. 2018).

### Phylogenetic Analyses and Age Estimates of MtDNA Haplogroups

The complete maximum parsimony (MP) tree was built by modifying a previously published mtPhyl v. 5.003 (Eltsov and Volodko 2011) script to account for new samples (Lombardo et al. 2022), haplogroups and control region mutations. Indels were not considered for tree construction. The Western house martin, *Delichon urbicum* (NC_050298) was used as an outgroup. Tree topology was confirmed also via a 1000 bootstrap MP tree in MEGAX v.11.0.11. Ages of haplogroups and sub-haplogroups were determined via molecular methods by performing Bayesian estimations using BEAST v.2.7.7 (Bouckaert et al. 2019) under the HKY (κ=2, estimated frequencies) nucleotide substitution model (8γ-distributed rates, 0.5 shape and estimated 0.5 proportion invariant sites) (Broggini et al. 2024) with a relaxed clock log-normal value of 0.0245 s/s/Myr (or one mutation every 2616 years) (Lombardo et al. 2022) . Current population size was set to a mean estimate (BirdLife International, 2019) value of 388 million individuals (uniform distribution, 487M upper, 290M lower). MCMC chain length was set to 50,000,000 iterations, with samples drawn every 1,000 steps after a burn-in of 10% using both log combiner and tree annotator (as in Ferrer Obiol et al., 2025). Bayesian skyline plots were constructed using median tree heights and 2000 bins using Tracer v.1.7.2 (Rambaut et al. 2018). Haplogroup expansion maps were built using Surfer v 19.1.189 (Golden Software, Inc., Golden, CO, USA, www.goldensoftware.com/) using the point kriging method (Figure 3) and overlayed on a world map.

## Author Contributions

Conceptualization, G.L. and G.B.; methodology, G.L.; software, G.L. with contributions from M.C. and A.DB.; formal analysis and investigation, G.L.; writing— original draft preparation, G.B., M.C. and G.L.; writing—review and editing, G.B., M.C., A.DB, G.R.G., L.G. and G.L. funding acquisition, G.B, L.G. All authors have read and agreed to the published version of the manuscript.

## Data Availability

The sequence data for the 181 mitogenomes are available in GenBank under accession numbers (in elaboration).

