## Supplementary Figures for "The end of the American dream: a hard to Swallow reality - How the Barn Swallow (*Hirundo rustica*) returned from America, a complete mtDNA phylogeny"

**Supplementary Material**

***Supplementary Figures***

**Figure S1. Detailed maximum parsimony phylogeny of *Hirundo rustica* mitogenomes.** This tree was built using the entire mitogenome coding-region and was rooted using *D. urbicum*. Main haplogroup and sub-haplogroup affiliations are shown. Sub-haplogroups were named only when encompassing at least two haplotypes. Subspecies affiliations are according to the colours in the legend. Mutations, relative to HrrRS, are transitions unless a base is explicitly indicated. Suffixes indicate transversions (to A, G, C, or T). Reversions are marked with "@" and recurring mutations are underlined. Both sample ID names (in green) and IDs employed in phylogenetic analyses are provided. Heteroplasmic positions are shown below sample names.

EXCEL FILE

Pre-print - Not Peer Reviewed

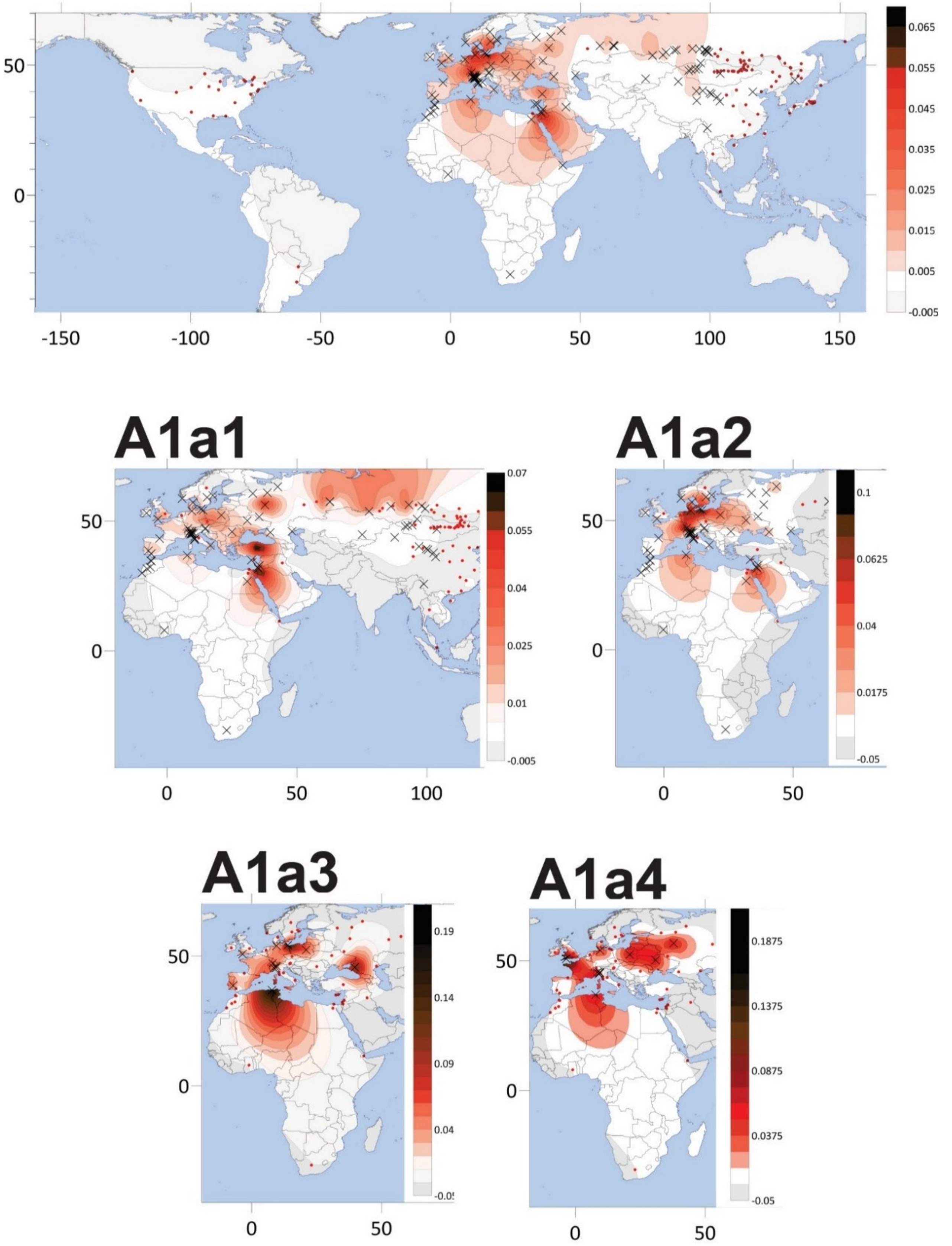


**Figure S2. Haplogroup frequency distribution maps of *Hirundo rustica rustica* main haplogroups.** Dots indicate the geographical locations of all sampled individuals; Crosses indicate individuals of subspecies in question. Colour scale indicates frequency of haplogroup in the given map.

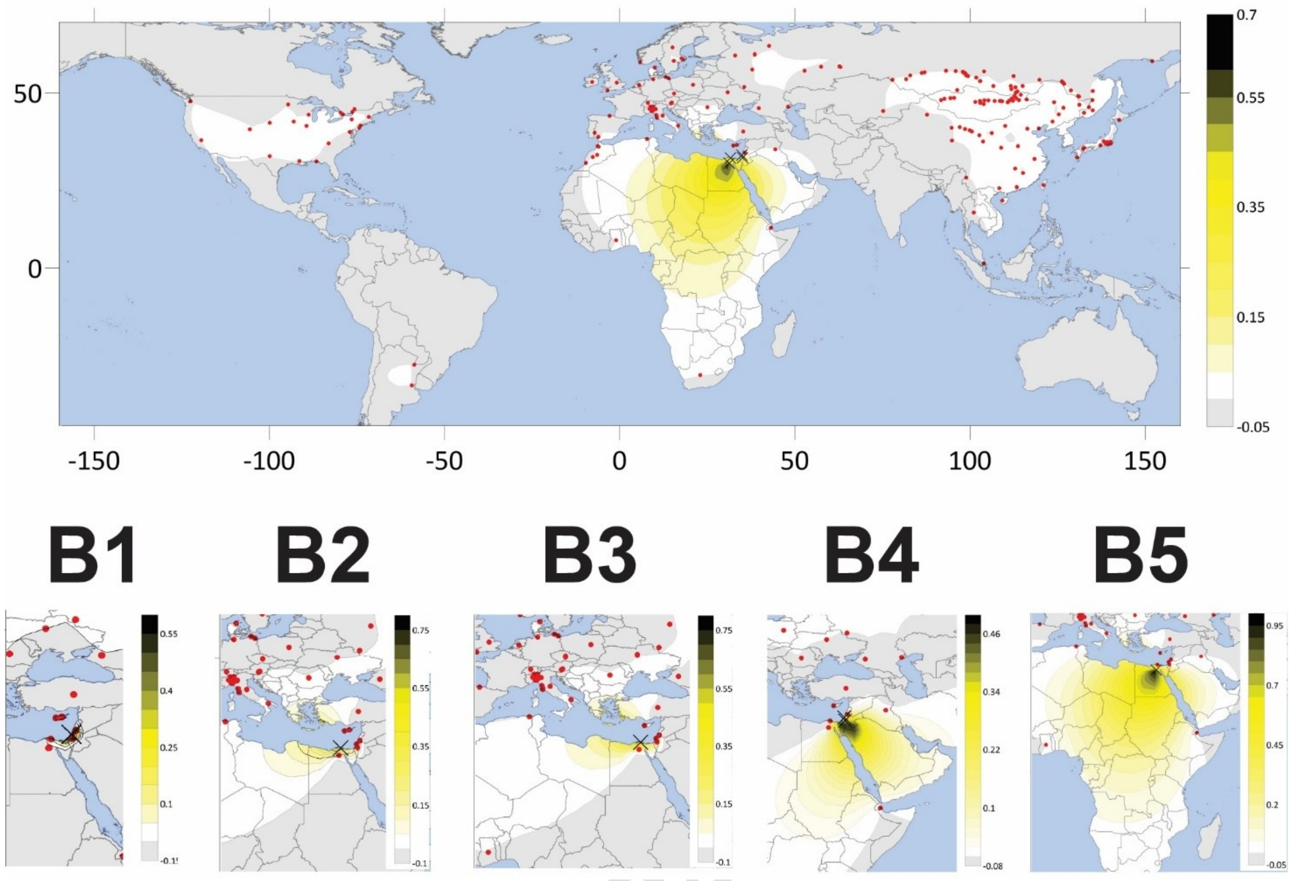


Pre-print - Not Pe

**Figure S3. Haplogroup frequency distribution maps of *Hirundo rustica savignii* main haplogroups.** Dots indicate the geographical locations of all sampled individuals; Crosses indicate individuals of subspecies in question. Colour scale indicates frequency of haplogroup in the given map.

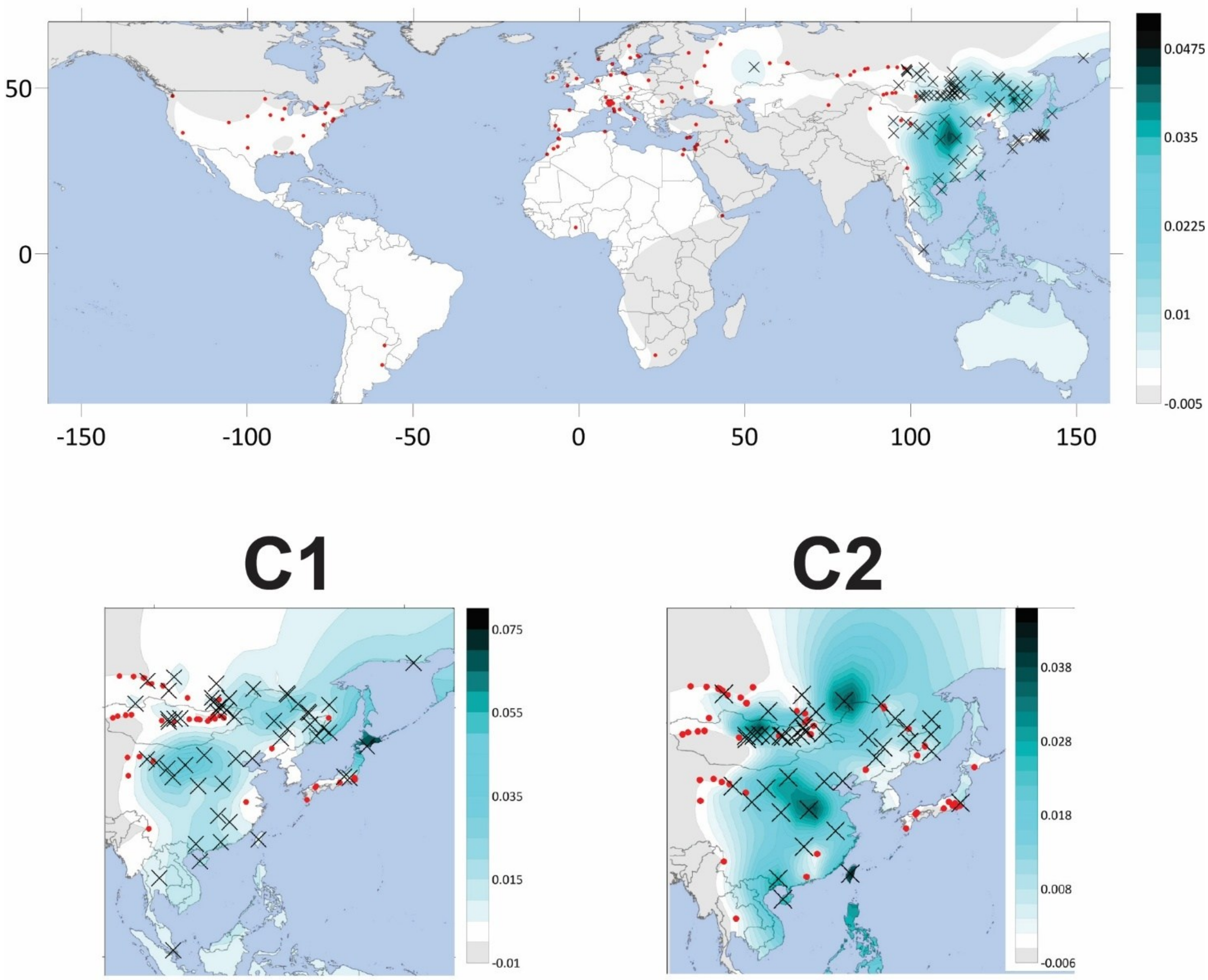


**Figure S4. Haplogroup frequency distribution maps of *Hirundo rustica gutturalis* main haplogroups.** Dots indicate the geographical locations of all sampled individuals; Crosses indicate individuals of subspecies in question. Colour scale indicates frequency of haplogroup in the given map.

Pre-print

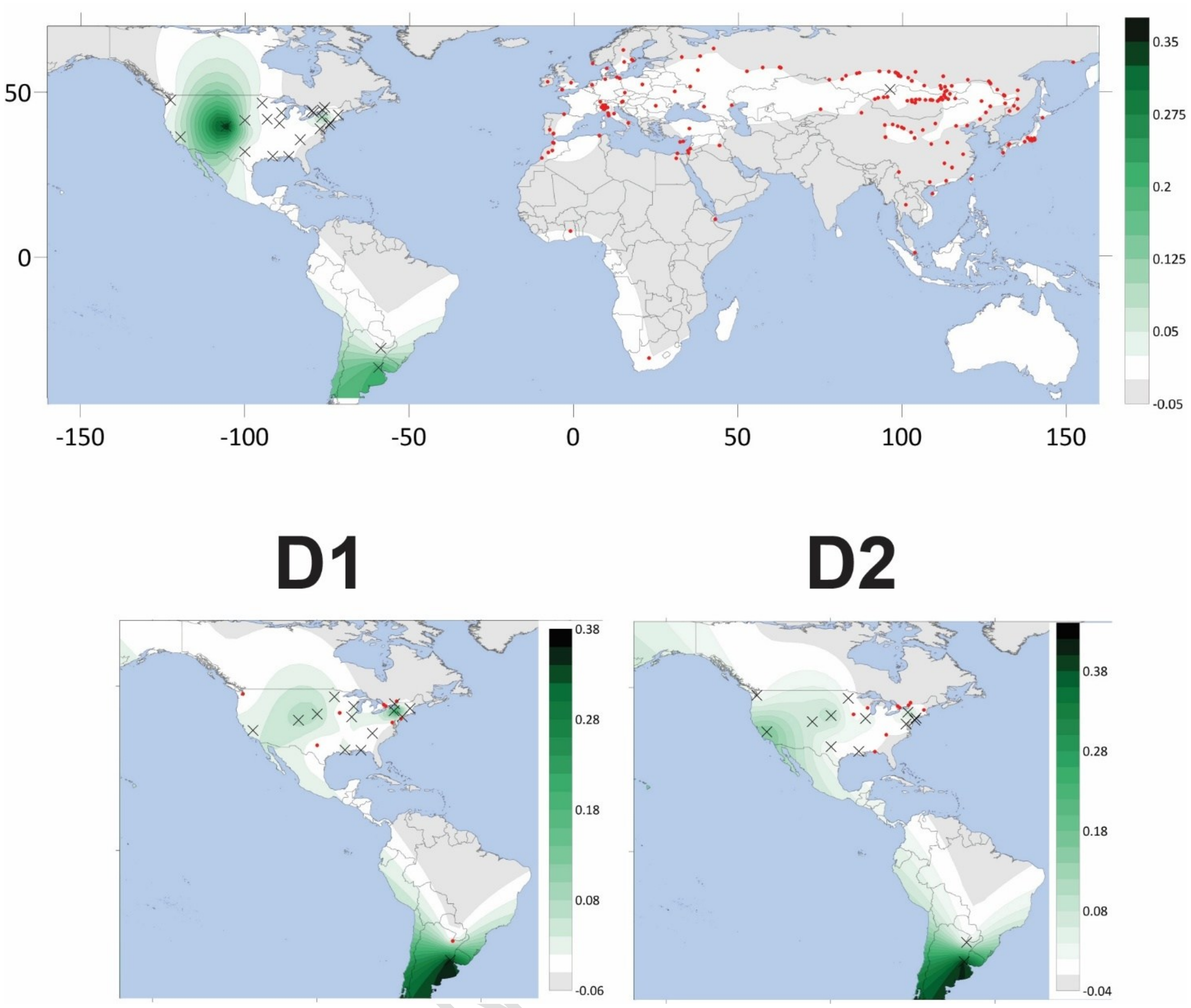


**Figure S5. Haplogroup frequency distribution maps of *Hirundo rustica erythrogaster* main haplogroups.** Dots indicate the geographical locations of all sampled individuals; Crosses indicate individuals of subspecies in question. Colour scale indicates frequency of haplogroup in the given map.

Pre-print

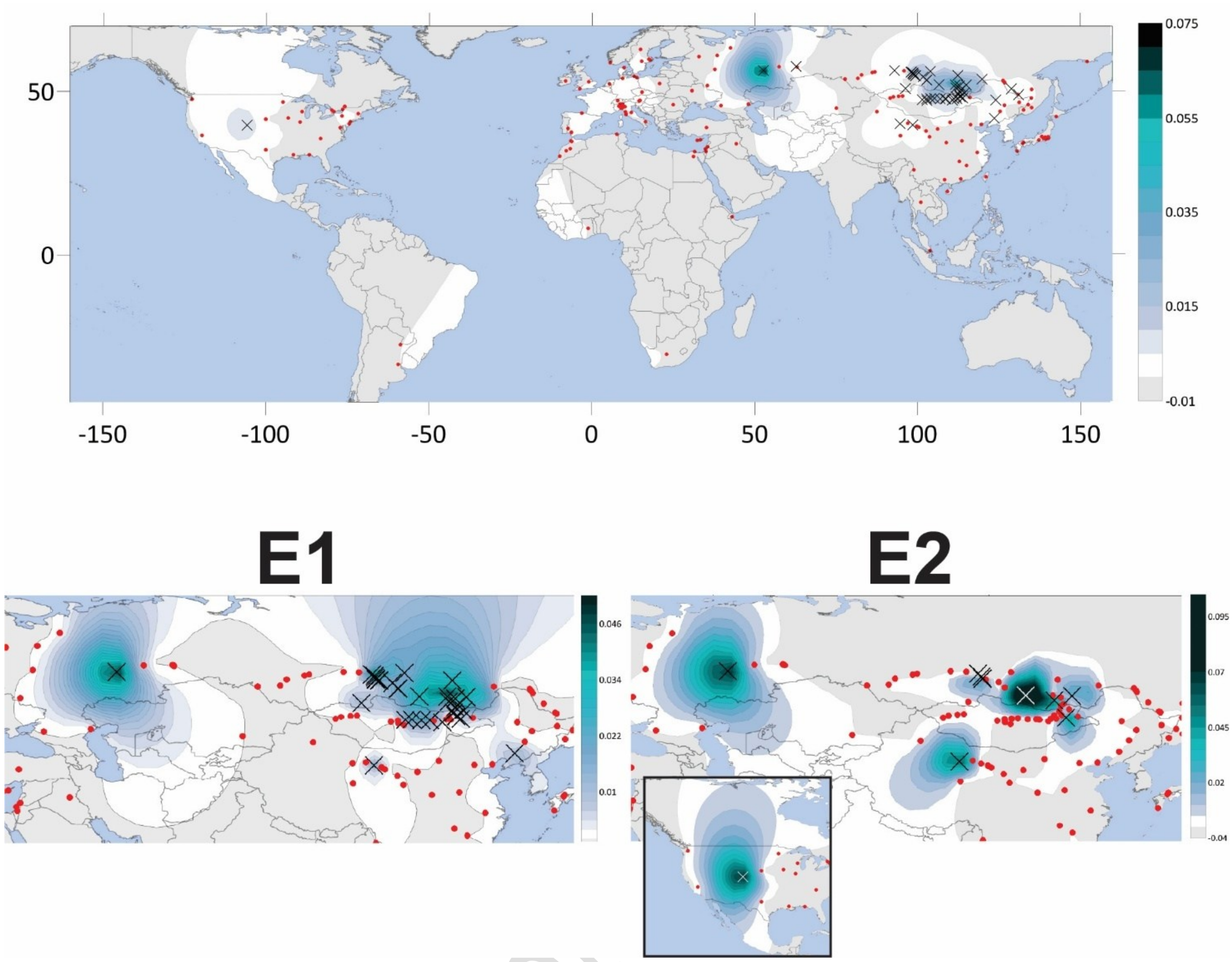


**Figure S6. Haplogroup frequency distribution maps of *Hirundo rustica tytleri* main haplogroups.** Dots indicate the geographical locations of all sampled individuals; Crosses indicate individuals of subspecies in question. Colour scale indicates frequency of haplogroup in the given map.

***Supplementary Tables***

**Table S1. Coalescence age estimates of haplogroups using cytb and whole mtDNA.** Bayesian age estimates for barn swallow haplogroups and sub-haplogroups and clade separation ages of all available species in the genus.

EXCEL FILE

Pre-print - No

Table S2. **Extended nucleotide diversity (%).** Within and between barn swallow main haplogroups with their respective SD. Shading goes from green, lowest diversity to red, highest. Intragroup nucleotide diversities (π) are on the diagonal.

| | A | B | C | D | E | HRAM | OUT | A HRR | A HRT | A1B | A2 | AB | CDE |
|---|---|---|---|---|---|---|---|---|---|---|---|---|---|
| A | 0.153 ±0.002 | 0.530 ±0.012 | 1.203 ±0.010 | 1.198 ±0.018 | 1.242 ±0.014 | 0.390 ±0.019 | 6.583 ±0.520 | 0.152 ±0.002 | 0.190 ±0.017 | 0.185 ±0.010 | 0.294 ±0.028 | 0.164 ±0.004 | 1.170 ±0.012 |
| B | | 0.086 ±0.007 | 1.186 ±0.056 | 1.190 ±0.096 | 1.237 ±0.079 | 0.426 ±0.025 | 6.605 ±0.812 | 0.556 ±0.014 | 0.520 ±0.037 | 0.558 ±0.117 | 0.594 ±0.148 | 0.542 ±0.013 | 1.215 ±0.062 |
| C | | | 0.159 ±0.006 | 0.706 ±0.024 | 0.730 ±0.020 | 1.020 ±0.017 | 6.592 ±1.171 | 1.200 ±0.044 | 1.198 ±0.120 | 1.201 ±0.413 | 1.196 ±0.473 | 1.200 ±0.041 | 0.748 ±0.069 |
| D | | | | 0.195 ±0.024 | 0.285 ±0.015 | 1.018 ±0.019 | 6.589 ±0.696 | 1.149 ±0.020 | 1.150 ±0.054 | 1.137 ±0.179 | 1.145 ±0.209 | 1.150 ±0.018 | 0.318 ±0.023 |
| E | | | | | 0.169 ±0.021 | 1.050 ±0.018 | 6.608 ±0.629 | 1.189 ±0.017 | 1.191 ±0.044 | 1.182 ±0.146 | 1.185 ±0.170 | 1.191 ±0.017 | 0.284 ±0.026 |
| HRAM | | | | | | 0.543 ±0.023 | 6.576 ±0.517 | 0.297 ±0.017 | 0.301 ±0.017 | 0.317 ±0.023 | 0.418 ±0.034 | 0.308 ±0.017 | 1.056 ±0.018 |
| OUT | | | | | | | 7.571 ±0.582 | 6.583 ±0.521 | 6.583 ±0.619 | 6.592 ±1.299 | 6.544 ±1.471 | 6.584 ±0.519 | 6.595 ±0.581 |
| A HRR | | | | | | | | 0.129 ±0.002 | 0.157 ±0.004 | 0.185 ±0.010 | 0.294 ±0.029 | 0.164 ±0.004 | 1.170 ±0.013 |
| A HRT | | | | | | | | | 0.191 ±0.024 | 0.186 ±0.024 | 0.292 ±0.048 | 0.169 ±0.006 | 1.171 ±0.033 |
| A1B | | | | | | | | | | 0.031 ±0.006 | 0.290 ±0.131 | 0.197 ±0.011 | 1.158 ±0.109 |
| A2 | | | | | | | | | | | 0.204 ±0.102 | 0.303 ±0.029 | 1.165 ±0.128 |
| AB | | | | | | | | | | | | 0.176 ±0.007 | 1.171 ±0.012 |
| CDE | | | | | | | | | | | | | 0.331 ±0.032 |

Pre-print - Not Peer Reviewed
